# Atomistic structure of the interlocking claw-like amyloid fibril of full-length glucagon

**DOI:** 10.1101/2022.11.21.517306

**Authors:** Hyeongseop Jeong, Yuxi Lin, Jin Hae Kim, Wookyung Yu, Yunseok Heo, Hyung-Sik Won, Masaki Okumura, Young-Ho Lee

## Abstract

Glucagon is a peptide hormone which posits a significant potential as a therapeutic molecule for various human diseases. One of the major challenges hampering medicinal application of glucagon, however, is its insoluble and aggregation-prone property. Although glucagon is dissolvable, it aggregates easily and forms amyloid fibrils. To date, despite many studies to understand how glucagon aggregates and fibrillizes, its detailed amyloidogenesis mechanism is still elusive, particularly due to insufficient structural information of glucagon amyloid fibrils. Here we report the novel structure of a glucagon amyloid fibril, which was determined with cryo-electron microscopy (cryo-EM) to a 3.8-Å resolution. Our model features with tight and extensive inter-monomer interactions, which efficiently stabilize the entire chain of full-length glucagon into the V-shape conformation, clamping the other monomer to construct a dimeric architecture orthogonal to the fibril axis. Notably, the current structure significantly differs from the previous solid-state NMR model, which gives direct evidence for multiple and dynamic amyloidogenesis mechanisms of glucagon. In addition, our results are expected to provide important insights to appreciate the molecular details of glucagon in its initial amyloidogenesis and fibril elongation processes.

## Introduction

Glucagon is a 29 amino-acid peptide hormone playing an essential role in regulating the blood glucose levels of humans.^1^ Although glucagon was first identified as an antagonistic molecule of insulin (thus, being named a *gluc*ose *agon*ist) in the 1920s,^2,3^ subsequent studies showed that its physiological importance is not limited to glucose metabolism. Glucagon is capable of regulating a wide range of physiological processes, such as lipid metabolism, amino acid metabolism, and energy expenditure, along with glucose level regulation.^4^ Of importance, this led to substantial expansion of the therapeutic potential of glucagon for various human diseases, not being limited to hypoglycemia but including dyslipidemia and obesity.^1,5^ However, one of the major challenges limiting therapeutic applications of glucagon is its aggregation-prone and amyloidogenic propensity.^6^ Although glucagon is soluble in a dilute condition, upon being in a more concentrated condition to ensure its therapeutic efficacy, glucagon becomes insoluble at neutral pH.^7,8^ Acidic or basic pH conditions could be made to increase the solubility; yet, a basic condition was shown to compromise its chemical integrity,^9^ whereas an acidic environment caused time-dependent amyloid formation of glucagon.^6,10^ These observations evidently point to the necessity of appreciating the aggregation and fibrillization mechanisms of glucagon for its wider and more appropriate applicability, which has been elusive particularly due to insufficient information regarding the structure of glucagon amyloid fibrils.

Structural studies of glucagon date back to 1970s, in which X-ray crystallographic and NMR-based studies reported its monomeric and highly flexible conformation in a dilute aqueous condition.^7,11^ Since then, numerous studies have been conducted to elucidate the structural features of soluble and insoluble glucagon in several conformational states by employing a wide range of techniques, *e.g*., X-ray crystallography,^11–14^ solution- and solid-state NMR,^8,15,16^ high-resolution atomic force microscopy,^17^ circular dichroism (CD),^10^ and FT-IR spectroscopy.^18^ All these works showed that the fibrillar aggregates of glucagon adopt β-strand-rich conformations as commonly found in amyloid fibrils.^10,15,18^ More recently, the first atomic-resolution model for glucagon amyloid fibrils was determined by solid-state NMR.^19^ This study demonstrated that the glucagon amyloid fibril accommodates two co-existing β-strand conformations which alternatively assemble into antiparallel β-sheets along the amyloid fibril axis, *i.e*., the cross-β structure. Most importantly, despite significant contribution of this work to advance our understandings for the glucagon aggregation mechanism, it is notable that structural polymorphism of glucagon amyloid fibrils was already reported.^17,20–22^ Thus, this indicates that more extensive research exploring diverse aggregation mechanisms and conformations of amyloid fibrils is necessary to fully resolve the aggregation-related issues of glucagon. In this work, we employed cryo-electron microscopy (cryo-EM) to acquire the atomic-resolution structure of glucagon amyloid fibrils that were assembled at acidic solution.

## Results

To investigate the structural details at atomic resolution, amyloid fibrils of full-length glucagon were first assembled at acidic solution (pH 2.0) including 100 mM NaCl. Real-time monitoring of thioflavin T (ThT) fluorescence at 37 °C showed a sigmoidal increase (Fig. 1a), indicating a nucleation-dependent amyloid fibrillation. The far-UV CD spectrum of glucagon before the incubation showed a single negative peak at ~200 nm (Fig. 1b), suggesting that the secondary structures of glucagon monomers are largely disordered. However, the CD spectrum of glucagon after incubation exhibited two negative peaks at ~205 and ~222 nm (Fig. 1b). This unusual CD spectrum was different from a typical CD spectrum of amyloid fibrils with a single minimum at ~218 nm, suggesting dynamic and polymorphic amyloid formation of glucagon. Indeed, distinct CD spectra have been observed for glucagon amyloid fibrils harboring different structures and morphologies.^20^ Our aggregation procedure successfully produced analyzable amyloid fibrils (Fig. 1c), from which the three-dimensional (3D) structure reconstitution analysis was conducted to have the high-resolution EM density map (Fig. 1d and Table 1).

**Figure 1.**
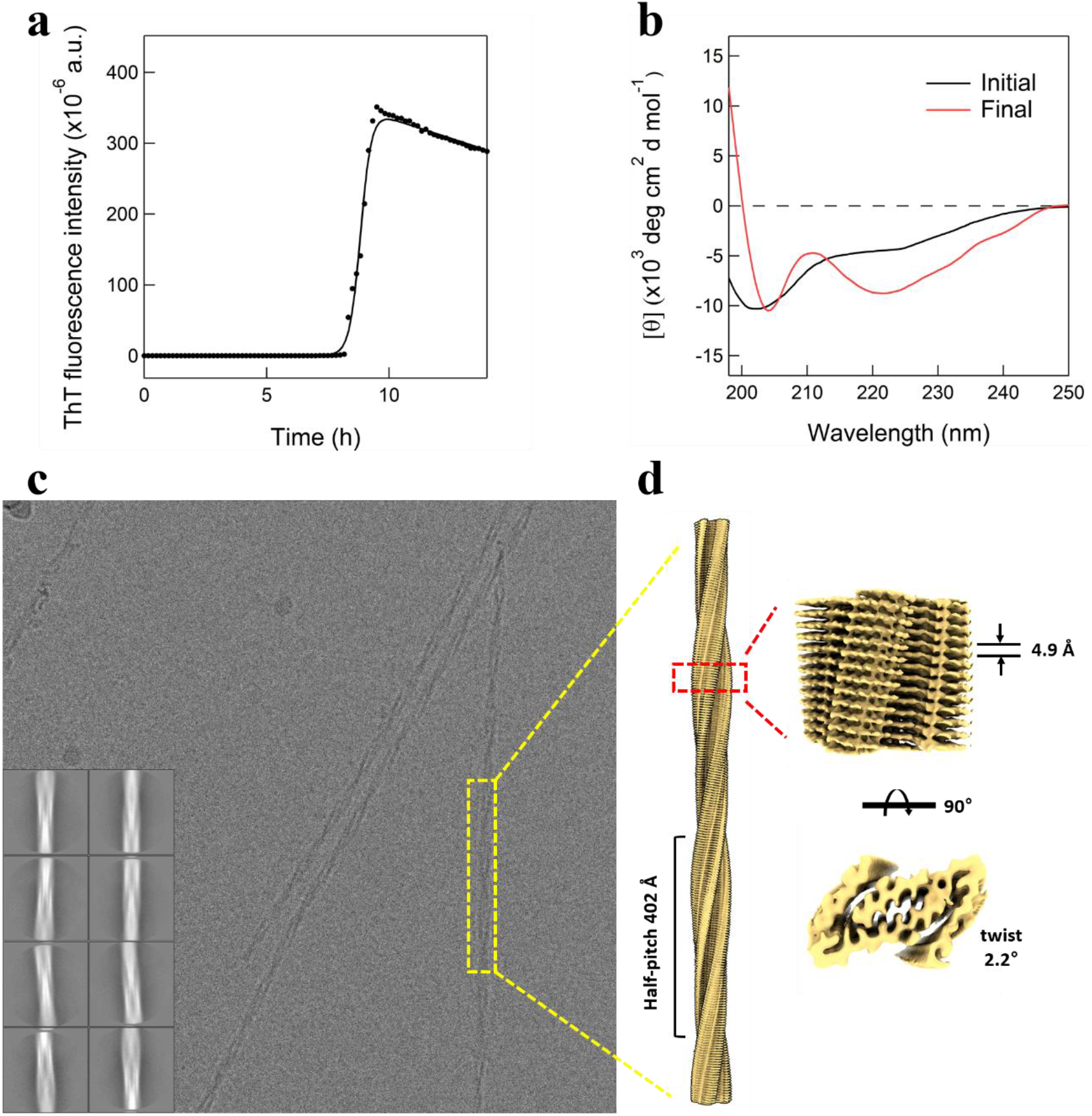
The formation of glucagon amyloid fibrils. (a) Kinetic profile of glucagon amyloid formation was traced using ThT fluorescence assay. The closed circles represent the raw data averaged from three separate samples. The solid line was drawn as an eye guide. (b) Far-UV CD spectra of glucagon before (black) and after (red) ThT fluorescence assay were acquired. The representative cryo-EM micrograph (c) and the electron density map (d) of glucagon amyloid fibrils showing their physical characteristics. Note that the glucagon amyloid fibril of the current study exhibits the right-handed helical assembly of glucagon dimers, which form the in-register cross-β conformation.

**Table 1.**
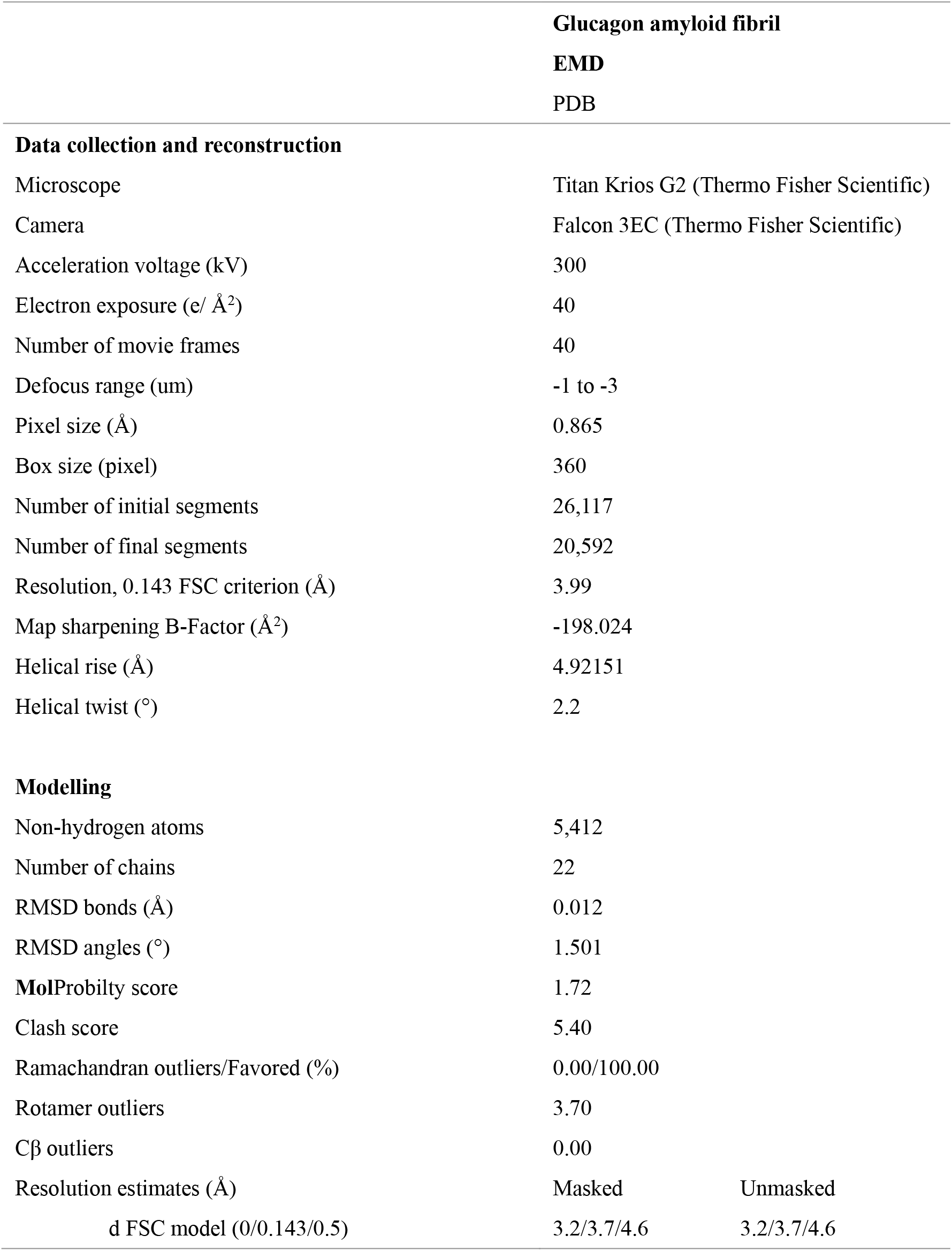
Cryo-EM data collection, reconstruction, and modelling parameters.

|  | Glucagon amyloid fibril |  |
| --- | --- | --- |
|  | EMD |  |
|  | PDB |  |
| **Data collection and reconstruction** |  |  |
| Microscope | Titan Krios G2 (Thermo Fisher Scientific) |  |
| Camera | Falcon 3EC (Thermo Fisher Scientific) |  |
| Acceleration voltage (kV) | 300 |  |
| Electron exposure (e/ Å²) | 40 |  |
| Number of movie frames | 40 |  |
| Defocus range (um) | -1 to -3 |  |
| Pixel size (Å) | 0.865 |  |
| Box size (pixel) | 360 |  |
| Number of initial segments | 26,117 |  |
| Number of final segments | 20,592 |  |
| Resolution, 0.143 FSC criterion (Å) | 3.99 |  |
| Map sharpening B-Factor (Å²) | -198.024 |  |
| Helical rise (Å) | 4.92151 |  |
| Helical twist (°) | 2.2 |  |
| **Modelling** |  |  |
| Non-hydrogen atoms | 5,412 |  |
| Number of chains | 22 |  |
| RMSD bonds (Å) | 0.012 |  |
| RMSD angles (°) | 1.501 |  |
| **Mol**Probilty score | 1.72 |  |
| Clash score | 5.40 |  |
| Ramachandran outliers/Favored (%) | 0.00/100.00 |  |
| Rotamer outliers | 3.70 |  |
| Cβ outliers | 0.00 |  |
| Resolution estimates (Å) | Masked | Unmasked |
| d FSC model (0/0.143/0.5) | 3.2/3.7/4.6 | 3.2/3.7/4.6 |

Based on this analysis, we determined the atomic-resolution model of the glucagon amyloid fibril and demonstrate its structural and physicochemical properties (Fig. 2). Our model indicated that a glucagon amyloid fibril is a right-handed helical assembly of glucagon dimers, which constitute parallel and in-register β-sheets. Remarkably, the new structural model is acknowledged by its unique feature that a glucagon monomer adopts a claw-like V-shape, grasping the other monomer to make a tight and extensive dimer interface (Fig. 2a). Almost all residues, including even the N- and C-termini of a glucagon monomer, contribute to construct this concrete architecture. In addition, the layer that is constituted by a glucagon dimer is not perfectly orthogonal to the fibril axis; it is tiled by about 10° with respect to the orthogonal plane of the fibril axis (Fig. 2b). These characteristics made each β-stand interlock with nearby five β-strands in a staggered fashion, fostering tight packing of amino acids.

**Figure 2.**
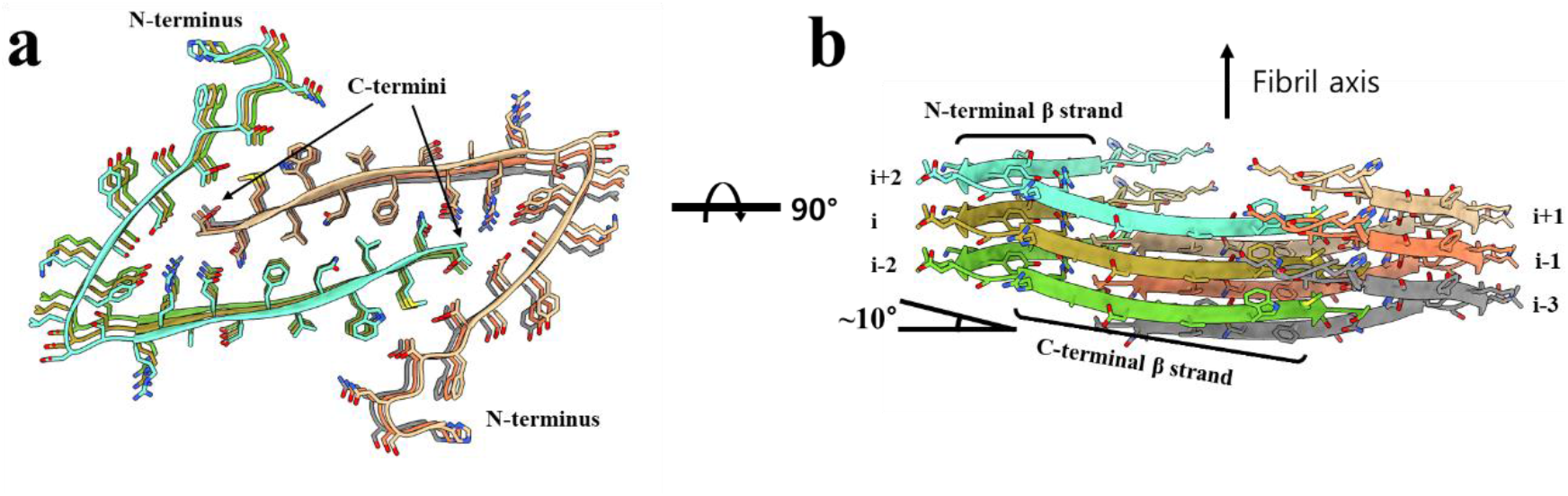
The cryo-EM structures of glucagon amyloid fibrils shown at the two different orientations. (a) Top view; the N-terminal and C-terminal β-strands of one glucagon monomer form a clamp-like conformation, which embraces the C-terminal β-strand from the other glucagon monomer. (b) Side view; the glucagon monomer makes ~10° tilted arrangement with respect to the fibril axis, constructing extensive inter-monomer interfaces with five monomeric subunits.

More specifically, there are a few notable features in the new structural model of a glucagon amyloid fibril. First, each glucagon monomer constructs two β-strands, the N-terminal (F6-L14) and the C-terminal β-strands (R17-N27). These two β-strands encompass the C-terminal β-strand of the other monomer at a dimeric architecture. Of note, the side chains of N- and C-terminal β-strands establish tightly packed inter-monomer interfaces, on which the side chain packing happens alternatively staggering with each other (Fig. 2b). For example, with respect to the fibril axis, the vertical position of the side chains from the C-terminal β-strand of the ‘i-1’ monomer locates between the side chains of the N- (upper) and C-terminal (lower) β-strands of the ‘i’ monomer (Fig. 2b). This staggered side-chain packing enables glucagon to make highly extensive inter-monomer interactions involving six glucagon molecules in total.

The inter-monomer interfaces of a glucagon dimer are mainly composed of hydrophobic residues along with a few polar residues (Fig. 3a). The hydrophobic residues, such as F22 and L26, form a firm and dry interface between the two C-terminal β-strands, which may work as a hydrophobic core stabilizing the overall architecture (Fig. 3b, c). On the other hand, the polar residues, Y13, R18, Q20, N28, and the C-terminal carboxylate group (T29) comprise the polar inter-monomer network, contributing not only to stability but also to structural preference toward the observed interlocking claw-like conformation of the glucagon amyloid fibril (Fig. 3b, d). This network is composed of the hydrogen bond between Q20 and N28 and the ionic interaction between the C-terminal carboxylate end and the guanidino group of R18. Interestingly, it appears that the V-shaped glucagon monomer is also supported by a few intra-monomer interactions. The residues L14-S16, which are at the bridge loop between the N- and C-terminal β-strands, are involved in a few intra-monomer hydrogen bonds; the hydroxyl group of S16 forms a hydrogen bond with the backbone carbonyl oxygen atom of L14. The hydroxyl group of Y13 and the guanidino group of R18 also form an intra-monomer hydrogen bond (Fig. 3b). These interaction network may strengthen and pose the turn structure between the N- and C-terminal β-strands. There are also notable intra-monomer interactions at the N-terminus, where the residues H1-F6 form a locally compact structure. The overall conformation of a glucagon monomer maintains a constrained state even at the N- and C-termini, as being consistently evidenced with the relatively uniform B-factor over the entire glucagon chain (Fig. 3e).

**Figure 3.**
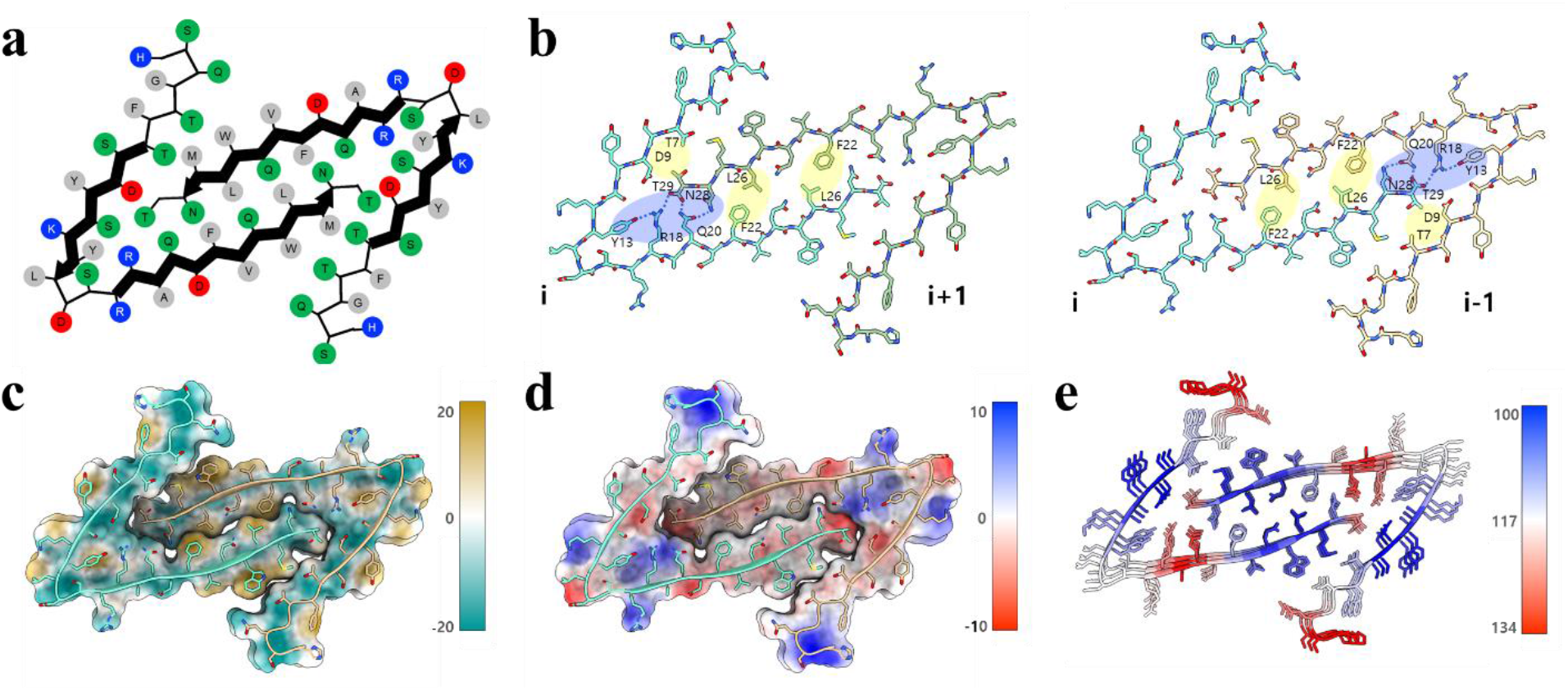
The amino-acid arrangement at the dimeric interfaces of glucagon amyloid fibrils. (a) The charge and polar properties along with their amino acid sequence are marked as a color; red, negatively charged residues; blue, positively charged residues; green, polar residues; grey, non-polar residues. Thick lines with an arrowhead indicate β-strands. (b) The representative inter- and intra-monomer interactions between amino acids are colored according to their polar nature; yellow, hydrophobic clusters; blue, polar networks. (c) The relative hydrophobic character is denoted on the surface; yellow, hydrophobic; green, hydrophilic. (d) The charge distribution is shown on the surface; red, negatively charged; blue, positively charged. (e) The cryo-EM model is colored according to their *B*-factor values. The B-factor scale is also shown on the right.

## Discussion

Owing to recent outstanding developments of cryo-EM and data analysis methodologies, a significant number of amyloid fibril structures have been determined at atomic resolution.^23^ Consequent boosts of available amyloid fibril structures allowed us to evidence various examples of structural polymorphisms that are originated from the same or similar proteins.^23^ Our cryo-EM structure of a glucagon amyloid fibril gives an excellent example to the polymorphic properties of amyloid fibrils at atomic resolution. To our knowledge, the only atomistic model of the amyloid fibril of glucagon was acquired by solid-state NMR, where relatively long and straight β-strand conformations were reported (Fig. 4).^19^ Remarkably, our cryo-EM analyses revealed the structure is significantly different from that of solid-state NMR, because the new model indicates that the two separate β-strands manifests within a monomer, and accommodates interlocking claw-like shape to embrace the C-terminal β-strand of the other monomer.

**Figure 4.**
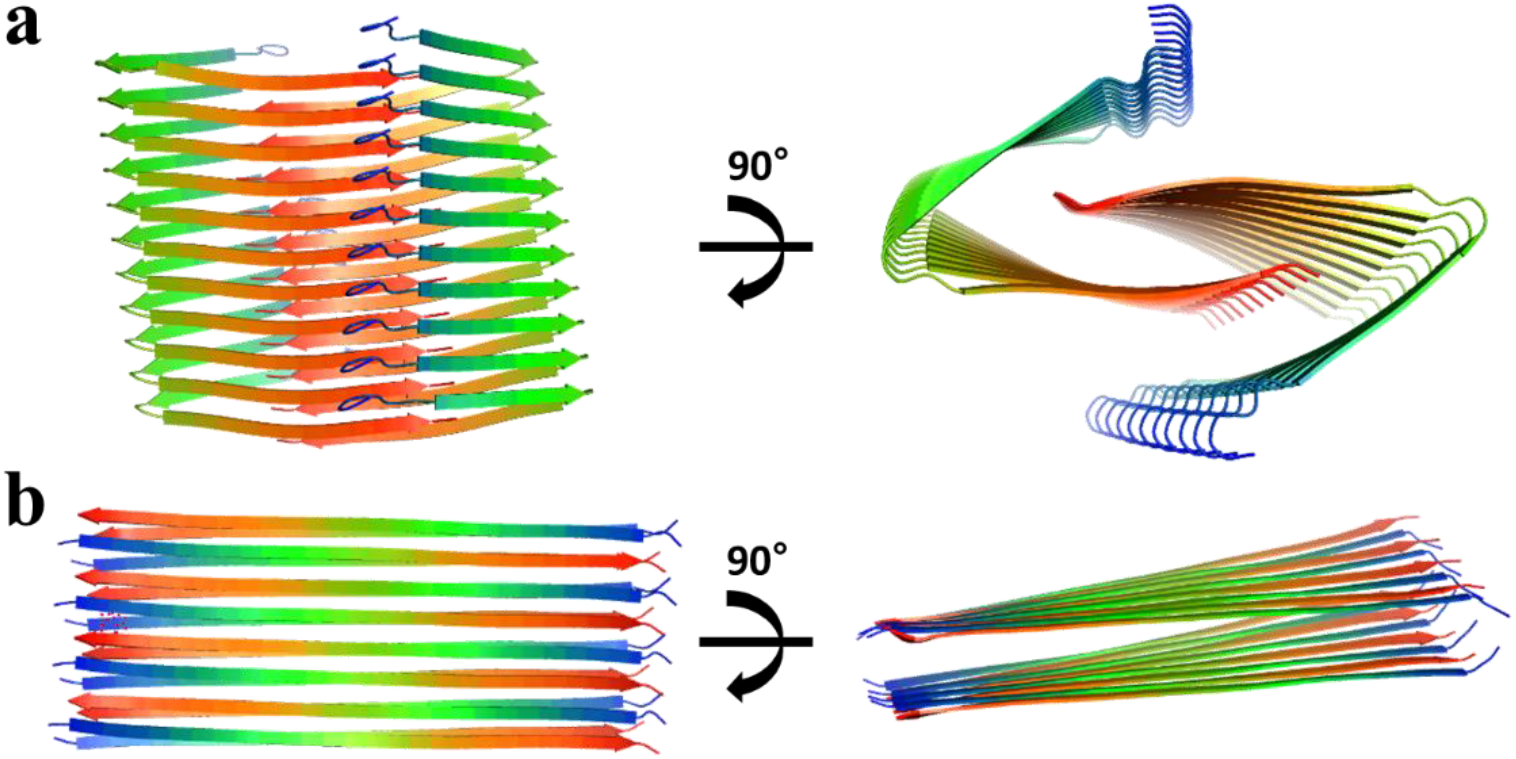
The structural models of polymorphic glucagon amyloid fibrils. (a) The structures determined in this study using cryo-electron microscopy, (b) The structural models revealed with solid-state NMR spectroscopy (PDB ID: 6nzn)^19^.

Previous solid-state NMR study used the amyloid fibrils which were made at the concentration of 8 mg ml^−1^ and 21 °C without a salt and shaking,^19^ whereas our amyloid fibrils were prepared at the concentration of 0.35 mg ml^−1^ and 37 °C in the presence of 100 mM NaCl and shaking. Indeed, a series of prior studies proved that the glucagon aggregation mechanism and the resultant morphology depend on the concentration of glucagon and the experimental condition such as temperature, pH, and ionic strength.^20,21,24–26^ Of note, it was reported that the high concentration of glucagon (*e.g*., 8 mg ml^−1^) facilitates the formation of relatively straight and extended protofilaments with a low stability, whereas glucagon in a dilute condition (*e.g*., 0.25 mg ml^−1^) fibrillizes into the twisted protofilaments that are relatively stable.^21,25^ Based on the observation that glucagon tends to form a trimer at a high concentration (*e.g*., > 1 mg ml^−1^), it was proposed that the trimeric intermediate state contributes to the formation of straight protofilament, while the monomeric state prefers to forming the twisted protofilament.^12,21^ Our data reported here are mostly aligned with these previous findings made in a dilute condition of glucagon.

Our interlocking claw-like structural model well explains the superior stability of the glucagon amyloid fibril that was formed in a dilute condition. Most importantly, the striking feature of our new model is that a glucagon monomer accommodates ~10° tilt with respect to the fibril axis, constructing highly extensive interfaces with five other glucagon monomers. The tilted layer of an individual monomer and the resultant expansion of the interaction with several monomers have been also observed in other amyloidogenic proteins, such as amyloid-β and islet amyloid polypeptide.^27,28^ However, the inter-monomeric interfaces of glucagon are even more extensive in that the N- and C-terminal β-stands of one monomer clamp the C-terminal β-strand of the other monomer not only on a two-dimensional plane but also along the fibril axis; the side chains of the N- and C-terminal β-stands makes a vertical clamp for the side chains of the C-terminal β-stand from another monomer. These observations raise an intriguing possibility that the N-terminal β-strand at the top of a glucagon amyloid fibril may remain flexible and work as a ‘prey’ for a subsequent monomer to be assembled. Upon catching the C-terminal region of a monomer, simultaneous stabilization of the N-terminal β-strand of the predecessor monomer and the C-terminal β-strand of the successor monomer may take place, thus efficiently elongating the amyloid fibril. Notably, this aggregation mechanism is more compatible with the monomeric state (thus being capable of accommodating a wider conformational space) than the trimeric (more constrained) state. On the one hand, despite its conformational rigidity, the trimeric state posits the N-terminus of one glucagon monomer at the proximity of the C-terminus of the other, which may contribute to efficient stacking of glucagon monomers in an alternative direction as in a straight yet less stable amyloid fibrils.

In addition, our structural model for the glucagon amyloid fibril gives considerable insights to devise the novel strategy of maximizing the therapeutic potential of glucagon and the related molecules while reducing their aggregation-prone and cytotoxic properties. In particular, a few previous studies reported that the variations in the C-terminus of glucagon, as seen from the glucagon and glucagon-like peptide-1 analogues whose C-terminus was amidated or extended with additional amino acids, were proven effective to increase therapeutic applicability and efficacy.^18,29,30^ Our structural model may help to provide a plausible scenario to explain these observations; the tight interaction network of the residues and the terminal carboxylic acid at the C-terminal region is critical to stabilize the amyloid conformation that was observed in the present study, and the perturbation of this interaction network results in repression of glucagon fibrillization. Taken together, we believe that our novel atomic-resolution model may help to design effective and easy-to-use glucagon-derived therapeutics as well as to elucidate the mechanisms of glucagon aggregation and the related possible cytotoxic processes.

## Methods

### Reagents

Glucagon (HSQGTFTSDYSKYLDSRRAQDFVQWLMNT; purity: >97%) was obtained from Toray Industries (Tokyo, Japan). Hydrochloric acid solution was purchased from Merck Millipore (Darmstadt, Germany). All other reagents were obtained from Sigma-Aldrich (Milwaukee, WI, USA).

### Preparation of glucagon amyloid fibrils

The stock solution of glucagon monomer was prepared as described previously.^25,26^ Briefly, the lyophilized glucagon was dissolved in the chilled, diluted HCl solution (pH 2.0) at approximately 0.55 mg ml^−1^. To remove undissolved aggregates, the solutions were subject to ultracentrifugation at 32,000 *g* for 15 min at 4 ^o^C. The upper half of the supernatant was collected, and the actual concentration of glucagon was determined by the absorbance and molar extinction coefficient (*ε*_280_ = 2.435 mL mg^−1^ cm^−1^) at 280. For the preparation of glucagon amyloid fibrils, the stock solution was further diluted to be 0.35 mg ml^−1^ of glucagon in the diluted HCl solution (pH 2.0) containing 100 mM NaCl. The 1^st^ generation (G_1_) amyloid fibrils were grown at 37 ^o^C in a 96-well microplate (Corning, Kennebunk, ME, USA) with continuous shaking for 1 day. The 2^nd^ generation (G2) amyloid fibrils were prepared by the addition of seeds (5% w/w) of G1 to a newly prepared glucagon monomer solution. Amyloid seeds were prepared by fragmenting long glucagon amyloid fibrils preformed with sonication.^31^ The seeded solution (G2) was incubated for 1 day with continuous shaking at 37 ^o^C. The same protocol of seeded amyloid formation was used for the subsequent generations, and the 4^th^ generation (G4) was finally used for Cryo-EM investigations.

### ThT fluorescence assay

The kinetics of glucagon amyloidogenesis was monitored using ThT fluorescence assay at 37 ^o^C. The sample solution (200 μL) of 0.35 mg ml^−1^ glucagon in the diluted HCl solution (pH 2.0) containing 100 mM NaCl and 5 μM ThT was loaded in triplicate into a 96-well microplate (Greiner-Bio-One, Tokyo, Japan). A sealing film (PowerSeal Cristal View, Greiner-Bio-One) was affixed to the microplate for preventing evaporation. The microplate was subject to the incubation with continuous shaking in a SpectraMax iD3 microplate reader (Molecular Devices, Sunnyvale, CA, USA). ThT fluorescence intensity was measured from the top of the microplate with the excitation and emission wavelengths of 445 and 490 nm, respectively.

### CD spectroscopy

Far-UV CD spectra of glucagon in the range from 198 to 250 nm were recorded using a JASCO J720 spectrophotometer (Tokyo, Japan) at room temperature. The sample containing 50 μM glucagon at pH 2.0 was used for CD measurements before and after ThT fluorescence assay. The scanning speed, the digital integration time, and the bandwidth were 100 nm/min, 2 s, and 1 nm, respectively. The light-path length of a CD cuvette was 1 mm. CD signals were expressed as the mean residue ellipticity (deg·cm^2^·dmol^−1^) after subtracting the background signal from the solution without glucagon.

### Specimen preparation for cryo-EM

Three microliter glucagon amyloid fibril sample (see above) was applied onto glow-discharged (15 mA current; negative charge; 60 sec) holey carbon grid (Quantifoil R1.2/1.3 Cu 200 mesh, Structure Probe, Inc., USA). The grid was blotted for 7 sec at 4°C with 100% humidity and vitrified using a Vitrobot Mark IV (Thermo Fisher Scientific, USA) by plunging into the liquid ethane cooled by liquid nitrogen.

### Cryo-EM data collection

Cryo-EM movies were collected using a transmission electron microscope (Titan Krios, Thermo Fisher Scientific, USA) equipped with a direct detector device camera (Falcon 3EC, Thermo Fisher Scientific, USA) with electron counting mode and automatic data acquisition software (EPU, Thermo Fisher Scientific, USA). Detailed data acquisition conditions and parameters are provided in Table 1.

### Image processing for 3D EM map and atomic model building

The dataset for glucagon amyloid fibril was processed using RELION 3.1-beta. The recorded movies were subjected to motion correction and CTF estimation using MotionCorr2 version 1.1.0 and ctffind version 4.1.10 respectively. Micrographs unsuitable for image processing such as those containing extremely low or high defocus and large motion drifts were removed. Then, 26,117 fibril segments were manually picked and extracted with an inter-box distance of ~10% and a box size of 360 × 360 pixels. Two rounds of reference-free 2D classification were performed using ~96% mask of the box size and 20,592 selected particles in ‘good’ aligned classes. To search the values of twist and rise, we conducted several rounds of 3D classification with K=4 and an initial model generated *de novo* from particles selected after 2D class averaging. The initial search values were used 4.6-5.1 (Å) for rise and 1.8-2.2 (deg) for twist which calculated from the crossover distance of dominant fibril morphology from raw micrographs and 2D classes. The refined 3D map was generated at 3.9 Å using the values 2.2 for twist and 4.92151 for rise those helical values were used by the calculated values that contributed for ‘good’ 3D map having the clearest ß-sheet (x-y plane) separation and peptide backbone from 3D classification. Then, the map was sharpened upon applying tight mask and using unfiltered half maps. The atomic model for glucagon amyloid fibrils was built manually in Coot version 0.9.1 and refined using phenix.real_space_refine in PHENIX version 1.19.2-4158. Detailed information for refinement and validation statistics was included in Table 1. Structure was visualized and figures were produced using UCSF ChimeraX version 1.3.

## Acknowledgements

This work was supported by the National Research Foundation of Korea (NRF) grant funded by the Korean government [NRF-2019R1A2C1004954 and NRF-2022R1A2C1011793 (Y.-H.L.) and NRF-2018R1C1B6008282 (J.H.K.)]; KBSI fund [C220000, C230130, and C280320] (Y.-H.L.), and National Research Council of Science & Technology (NST) grant funded by the Korean government [CCL22061-100 (Y.-H.L.)].

## References

1. Habegger, K. M. et al. The metabolic actions of glucagon revisited. Nat. Rev. Endocrinol. 6, 689–697 (2010).

2. Murlin, J. R., Clough, H. D., Gibbs, C. B. F. & Stokes, A. M. Aqueous Extracts of Pancreas. J. Biol. Chem. 56, 253–296 (1923).

3. Robison, G. A., Butcher, R. W. & Sutherland, E. W. Cyclic AMP. Annu. Rev. Biochem. 37, 149–174 (1968).

4. Janah, L. et al. Glucagon receptor signaling and glucagon resistance. Int. J. Mol. Sci. 20, (2019).

5. Day, J. W. et al. A new glucagon and GLP-1 co-agonist eliminates obesity in rodents. Nat. Chem. Biol. 5, 749–757 (2009).

6. Beaven, G. H., Gratzer, W. B. & Davies, H. G. Formation and Structure of Gels and Fibrils from Glucagon. Eur. J. Biochem. 11, 37–42 (1969).

7. Boesch, C., Bundi, A., Oppliger, M. & Wüthrich, K. ^1^H Nuclear-Magnetic-Resonance Studies of the Molecular Conformation of Monomeric Glucagon in Aqueous Solution. Eur. J. Biochem. 91, 209–214 (1978).

8. Braun, W., Wider, G., Lee, K. H. & Wüthrich, K. Conformation of glucagon in a lipid-water interphase by 1H nuclear magnetic resonance. J. Mol. Biol. 169, 921–948 (1983).

9. Caputo, N. et al. Mechanisms of glucagon degradation at alkaline pH. Peptides 45, 40–47 (2013).

10. Onoue, S. et al. Mishandling of the Therapeutic Peptide Glucagon Generates Cytotoxic Amyloidogenic Fibrils. Pharm. Res. 21, 1274–1283 (2004).

11. Sasaki, K., Dockerill, S., Adamiak, D. A., Tickle, I. J. & Blundell, T. X-ray analysis of glucagon and its relationship to receptor binding. Nature 257, 751–757 (1975).

12. Sturm, N. S. et al. Structure-function studies on positions 17, 18, and 21 replacement analogues of glucagon: The importance of charged residues and salt bridges in glucagon biological activity. J. Med. Chem. 41, 2693–2700 (1998).

13. Siu, F. Y. et al. Structure of the human glucagon class B G-protein-coupled receptor. Nature 499, 444–449 (2013).

14. Zhang, H. et al. Structure of the glucagon receptor in complex with a glucagon analogue. Nature 553, 106–110 (2018).

15. Yamane, I. et al. Fibrillation mechanism of glucagon in the presence of phospholipid bilayers as revealed by ^13^C solid-state NMR spectroscopy. Chem. Phys. Lipids 219, 36–44 (2019).

16. Haya, K., Makino, Y., Kikuchi-Kinoshita, A., Kawamura, I. & Naito, A. ^31^P and ^13^C solid-state NMR analysis of morphological changes of phospholipid bilayers containing glucagon during fibril formation of glucagon under neutral condition. Biochim. Biophys. Acta - Biomembr. 1862, 183290 (2020).

17. De Jong, K. L., Incledon, B., Yip, C. M. & DeFelippis, M. R. Amyloid fibrils of glucagon characterized by high-resolution atomic force microscopy. Biophys. J. 91, 1905–1914 (2006).

18. Onoue, S. et al. Structural transition of glucagon in the concentrated solution observed by electrophoretic and spectroscopic techniques. J. Chromatogr. A 1109, 167–173 (2006).

19. Gelenter, M. D. et al. The peptide hormone glucagon forms amyloid fibrils with two coexisting β-strand conformations. Nat. Struct. Mol. Biol. 26, 592–598 (2019).

20. Pedersen, J. S. et al. The changing face of glucagon fibrillation: Structural polymorphism and conformational imprinting. J. Mol. Biol. 355, 501–523 (2006).

21. Andersen, C. B., Otzen, D., Christiansen, G. & Rischel, C. Glucagon amyloid-like fibril morphology is selected via morphology-dependent growth inhibition. Biochemistry 46, 7314–7324 (2007).

22. Pedersen, J. S., Andersen, C. B. & Otzen, D. E. Amyloid structure - one but not the same: The many levels of fibrillar polymorphism. FEBS J. 277, 4591–4601 (2010).

23. Sawaya, M. R., Hughes, M. P., Rodriguez, J. A., Riek, R. & Eisenberg, D. S. The expanding amyloid family: Structure, stability, function, and pathogenesis. Cell 184, 4857–4873 (2021).

24. Jeppesen, M. D., Hein, K., Nissen, P., Westh, P. & Otzen, D. E. A thermodynamic analysis of fibrillar polymorphism. Biophys. Chem. 149, 40–46 (2010).

25. Andersen, C. B. et al. Glucagon fibril polymorphism reflects differences in protofilament backbone structure. J. Mol. Biol. 397, 932–946 (2010).

26. Ghodke, S. et al. Mapping out the multistage fibrillation of glucagon. FEBS J. 279, 752–765 (2012).

27. Gremer, L. et al. Fibril structure of amyloid-β (1–42) by cryo–electron microscopy. Science (80-.). 119, 116–119 (2017).

28. Röder, C. et al. Cryo-EM structure of islet amyloid polypeptide fibrils reveals similarities with amyloid-β fibrils. Nat. Struct. Mol. Biol. 27, 660–667 (2020).

29. Chabenne, J. R., DiMarchi, M. A., Gelfanov, V. M. & DiMarchi, R. D. Optimization of the native glucagon sequence for medicinal purposes. J. Diabetes Sci. Technol. 4, 1322–1331 (2010).

30. Lima, L. M. T. R. & Icart, L. P. Amyloidogenicity of peptides targeting diabetes and obesity. Colloids Surfaces B Biointerfaces 209, (2022).

31. Ikenoue, T. et al. Cold denaturation of α-synuclein amyloid fibrils. Angew. Chemie - Int. Ed. 53, 7799–7804 (2014).

